# Genome Wide Meta-Analysis identifies new loci associated with cardiac phenotypes and uncovers a common genetic signature shared by heart function and Alzheimer’s disease

**DOI:** 10.1101/386680

**Authors:** MªEugenia Sáez, Antonio González-Pérez, Begoña Hernández-Olasagarre, Aida Beà, Sonia Moreno-Grau, Itziar de Rojas, Gemma Monté, Adela Orellana, Sergi Valero, Joan X. Comella, Daniel Sanchis, Agustín Ruiz, For the Alzheimer’s Disease Neuroimaging Initiative

## Abstract

**Aims:** Echocardiography has become an indispensable tool for the study of heart performance, improving the monitoring of individuals with cardiac diseases. Diverse genetic factors associated with echocardiographic measures of heart structure and functions have been previously reported. The impact of several apoptotic genes in heart development identified in experimental models prompted us to assess their potential association with indicators of human cardiac function. This study started with the aim to investigate the possible association of variants of apoptotic genes with echocardiographic traits and to identify new genetic markers associated with cardiac function.

**Methods and results:** Genome wide data from different studies were obtained from public repositories. After quality control and imputation, association analyses confirm the role of caspases and other apoptosis related genes with cardiac phenotypes. Moreover, enrichment analysis showed an over-representation of genes, including some apoptotic regulators, associated with Alzheimer’s disease (AD). We further explored this unexpected observation which was confirmed by genetic correlation analyses.

**Conclusions:** Our findings show the association of apoptotic gene variants with echocardiographic indicators of heart function and reveal a novel potential genetic link between echocardiographic measures in healthy populations and cognitive decline later on in life. These findings may have important implications for preventative strategies combating Alzheimer’s disease.

## INTRODUCTION

Echocardiographic assessment of cardiac structure offers prognostic information about cardiac conditions such as heart failure (HF).^1^ Pathological processes including cardiomyocyte cell death, inflammatory cell response and changes in interstitial tissue of the heart are factors leading to adverse remodelling and HF.^2^

A number of apoptotic genes have been investigated as potential targets to prevent cardiomyocyte death, but it is now increasingly evident that caspase-dependent cell death plays a minor if any role in adult myocyte loss,^3^ which involves Cyclophilin D^4^ and calpains.^5^ By contrast, caspase proteins are now recognized as important factors for initial differentiation of stem cells to cardiomyocytes^6^ and its deficiency in vivo was shown to induce abnormal heart development.^7,8^ In rodent cardiomyocytes, caspase-3 is involved in WNT signalling and myocyte growth^9,10^ and also contributes to muscle-specific gene splicing by cleaving PTB.^11^ In addition, apoptotic DNA nucleases were shown to play a role in the developmental process of *C.elegans* including the *C.elegans* Caspase-associated DNase (CAD), ENDOG and TATD orthologs.^12^ Furthermore, ENDOG also contributes to the signalling pathways determining myocyte size^13^ through the control of reactive oxygen radicals (ROS).^14^ These facts lead us to hypothesize that caspases and the nucleases ENDOG and TATD play relevant functions in cardiomyocyte proliferation and maturation during development.

Genome wide association studies have been performed for evaluating comprehensive sets of echocardiographic traits in well characterized individuals included in large cohort studies.^15,16^ Using data from publicly available repositories, we aimed to explore the association between a selected group of candidate apoptosis-related genes and these echocardiographic phenotypes by means of meta-GWAS. Furthermore, we aimed to assess previously reported signals in our study datasets and performed an agnostic analysis to investigate relevant pathways revealed for each trait. Following the leading results of this analysis, we further explored the unsuspected genetic relationship between Alzheimer’s Disease (AD) and these echocardiographic traits, by estimating their genetic correlation, and identifying common genetic determinants of these conditions.

## METHODS

### STUDY COHORTS

#### Heart study cohorts

The four cardiovascular datasets analysed in this study were downloaded from dbGAP (https://www.ncbi.nlm.nih.gov/gap) after requesting the appropriate permissions. In the case of a multi-ethnic study, only Caucasian samples after principal component analysis (PCA) were retained for analysis. A summary of the clinical characteristics of these populations is shown in Table 1. A full description of each of them is provided as a Supplementary Note.

#### Alzheimer’s disease study cohorts

A total of seven AD datasets were used to further explore the observed enrichment of top genes derived from the analysis of echocardiographic traits on genes involved in AD pathways. As for previously described datasets, only Caucasian samples after principal component analysis (PCA) were retained for analysis. Demographic characteristics of these datasets are summarized in the Supplementary Table 1 whereas a full description is provided as a Supplemental Note.

### PHENOTYPES

#### Echocardiographic Phenotypes

Data from the most recent available echocardiographic examinations of each cohort were included in this study. The following five phenotypes were analysed: Left Ventricular Mass (LVM) (g), End-Diastolic Diameter of the Aortic Root (AROT) (cm), End-Diastolic LV Internal Dimension (LVID) (cm), Left Atrial Size (LAS) (cm), and LV Wall Thickness (LVWT)(cm). The latter was defined as the sum of the End-Diastolic Thicknesses of the Posterior Wall (TPW) and End-Diastolic Thicknesses of the Interventricular Septum (TIS). LVM was calculated using the formula 0.8 [1.04{(LVID + TIS + TPW)^3^ −(LVID)^3^}] + 0.6.^17^

### GENOTYPING AND IMPUTATION

The cardiovascular datasets included in this study were genotyped using different platforms: CARDIA and MESA were genotyped using the Affymetrix Genome-Wide Human 6.0 array, whereas the FHS was genotyped using the Affymetrix Human 500k array and the CHS cohort with the Illumina HumanCNV370-Duo v1.0.

AD datasets were genotyped using the Illumina arrays Human 610-Quad BeadChip (ADNI1, AddNeuroMed batch 1), the HumanOmniExpress BeadChip (ADNI2/GO, AddNeuroMed batch 2, ADGC dataset 3), the Human660W-Quad (ADGC datasets 1&2) and the HumanHap300-Duo BeadChip (The Mayo study) or the Affymetrix 250k NspI (the Neocodex-Murcia study), 500k (the TGEN and GenADA studies) or 6.0 (ROSMAP study) arrays.

Prior to imputation, we first performed an extensive quality control excluding individuals with more than 3% missing genotypes, with excess autosomal heterozygosity (>0.35), those showing a discrepancy between genotypic and reported sex, as well as individuals of non-European ancestry based on PCA analyses using SMARTPCA.^18^ At the genotype level, we removed SNPs with missing genotype rate > 5%, not in Hardy-Weinberg equilibrium (p<10^-6^ in controls) and SNPs with minor allele frequency (MAF) < 1%. Duplicated and related individuals were identified and removed by means of IBS estimates within and across studies.

Genotype imputation was performed using the minimac3 algorithm and the SHAPEIT tool for haplotype phasing at the University of Michigan server using the HRC reference panel (Das et al. 2016). After imputation, SNPs with an R2 quality estimate lower than 0.3 were excluded from further analyses.

### STATISTICAL ANALYSIS

All analyses were performed in Caucasian populations only. Individuals with prevalent myocardial infarction (MI) or congestive heart failure (CHF) were excluded from the study. Linear regression models available from PLINK software^19^ were fitted to investigate the association between genotypes and quantitative phenotypes, with age, sex, body mass index and the four principal components as covariates. For each phenotype, we obtained summary estimates of association across studies by using a fixed-effects model meta-analysis procedure implemented also in PLINK. For the genome wide SNP analysis, the conventional GWAS significance threshold was used (p=5 x 10^-8^).^20^

Gene-wise statistics were computed using MAGMA software, which takes into account physical distance and linkage disequilibrium (LD) between markers.^21^ Only SNPs with MAF above 1% were used in these analyses. At each trait, genes were ranked according to the global p mean value. For the candidate gene analysis, considering the number of traits (n=5) and genes (n=20) being explored, we set the threshold for study-wide statistical significance in 5 x 10^-4^. The top 200 genes from each association analysis were further explored for enrichment in known pathways using the R packages Webgestalt and enrichR^22,23^ with default parameters. The threshold for statistical significance in these analyses was FDR values below 0.1.

In order to explore genetic correlation between different traits we used a bivariate GREML analysis with GTCA software^24^ with default parameters. Furthermore, we obtained summary estimates of association across phenotypes performing unweighted meta-analysis of Fisher p-values. The threshold for statistical significance in all these analyses was p-values below 0.05.

## RESULTS

Overall, our study included data from 11,559 individuals with echocardiographic phenotypes from four different datasets (Table 1). After imputation and quality control, we obtained about 7 million SNPs with MAF>0.01 that were tested for association with echocardiographic traits at each study. We then performed a meta-GWAS to obtain summary estimates of association for each SNP. Genomic inflation factor (λ) ranged from 0.994 to 1.022 in these analyses, indicating absence of population stratification due to hidden population structure (Figure 1). MAGMA software was used for summarising the meta-GWAS SNP results in order to obtain a gene-wise statistic of the association between 18,480 genes and the five phenotypes.

**Figure 1.**
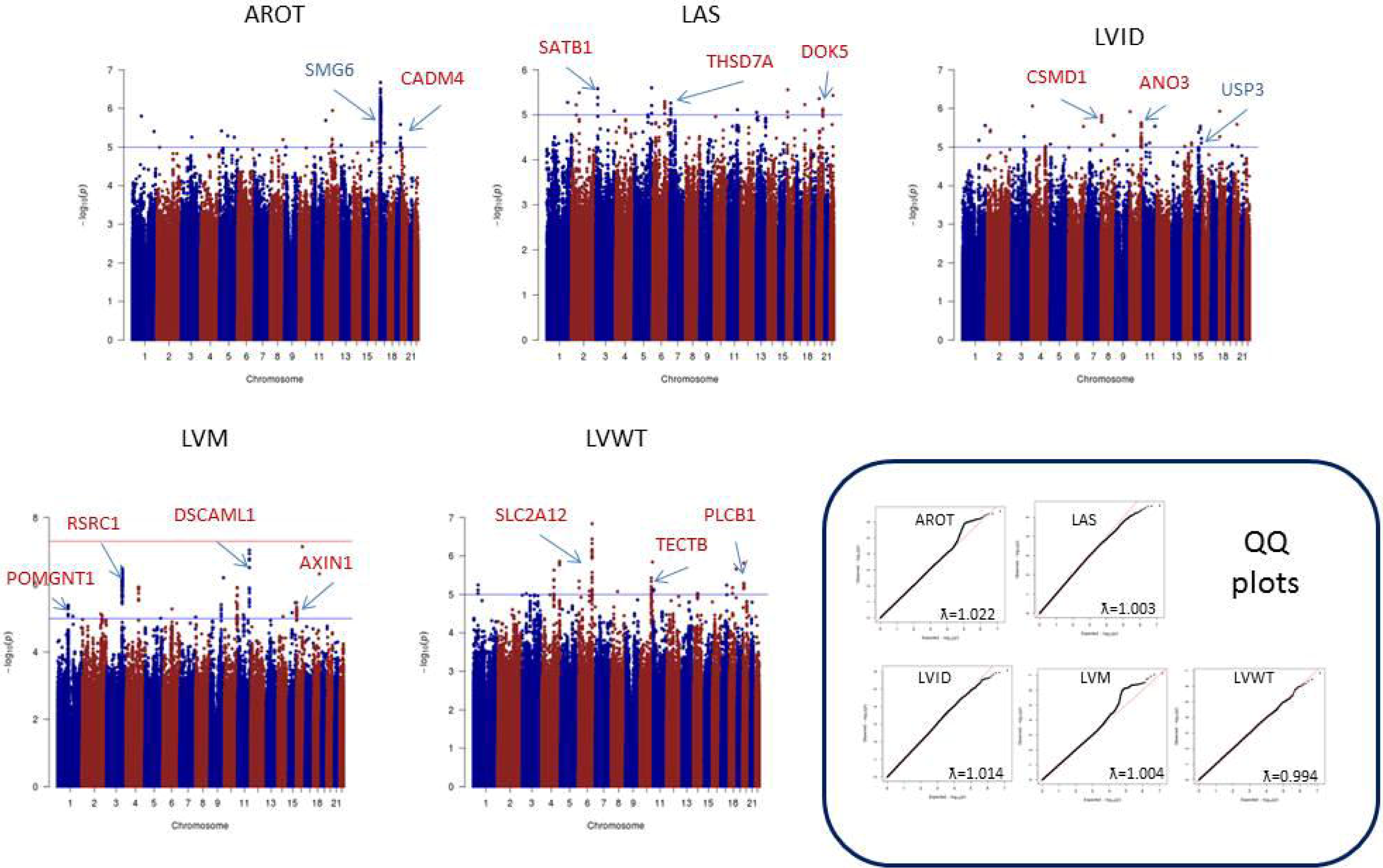
Manhattan plots of the meta-analyses for the different echocardiographic traits analysed. The threshold for genome-wide significance (P < 5 x 10-8) is indicated by the red line, while the blue line represents the suggestive threshold (P < 1 x 10-5). Loci previously associated with echocardiographic traits are shown in blue, and newly associated loci are shown in red.

### Association of apoptosis-related genes with cardiac phenotypes

Because there is experimental evidence supporting the role of some apoptosis-related genes with cardiac development and disease, we first analysed the potential association of polymorphisms in a series of apoptosis-related genes with the cardiac phenotypes. Table 2 shows the association results of the 20 apoptosis-related candidate genes. Study-wide statistically significant results were observed for the association of a genetic locus on 2q33.1 involving two initiator caspases (*CASP8* and *CASP10*) and the apoptosis regulator protein CFLAR (*CASP8 And FADD Like Apoptosis Regulator, c-FLIP*) with LVM. The same three genes were also linked to LVID with suggestive results (p<0.05), along with the Fas receptor-associated adaptor *FADD* (Fas-associated protein with death domain) and *BCL2* (B-cell lymphoma-2). *BCL2*, *FADD* and *TATDN1* (TatD DNase domain containing-1) showed suggestive signals of association with AROT. We did not find evidence of association of any calpain family member with any of the analysed traits (data not shown).

### Agnostic GWAS of genetic variants associated with cardiac phenotypes and enrichment analysis of top genes on echocardiographic traits

Although we did not find any GWAS significant signal at SNP-level (p<5x10^-8^) related with the analysed phenotypes, we observed several suggestive signals at the p<10^-5^ level (Supplementary tables 2-6), most of them intragenic (Figure 1). For each genotype, we ranked genes according to the MAGMA computed SNP-wise p-value (Table 3 and Supplementary tables 7-11).

Top 200 ranked genes were tested for enrichment using Webgestalt and EnrichR R packages (Supplementary tables 12-16). These top genes show little overlap across phenotypes (Figure 2a), which is reflected in the little overlap of the top ten enriched gene ontology (GO) categories (Figure 2b). For both AROT and LVID, we found enrichment on members from the keratin II gene cluster. LVID and LVWT share genes from the olfactory receptor families 52 and 56, whereas LVM and LVWT show enrichment on genes from the haemoglobin locus. Specific signals for LVID arise from genes involved in long QT signals and atrial fibrillation, the *HLA* locus and the vesicular and lysosomal system for LAS and grow factors such as *VEGF* and *FGF* were observed for LVWT. Surprisingly, in pathway analysis, we observed an enrichment in Alzheimer related pathways for both LVID and LVM, involving the *PSEN2*, *MPO*, *SMAD1*, *ADRB2*, *APH1B*, *MAPK9* and *RBP* genes.

**Figure 2.**
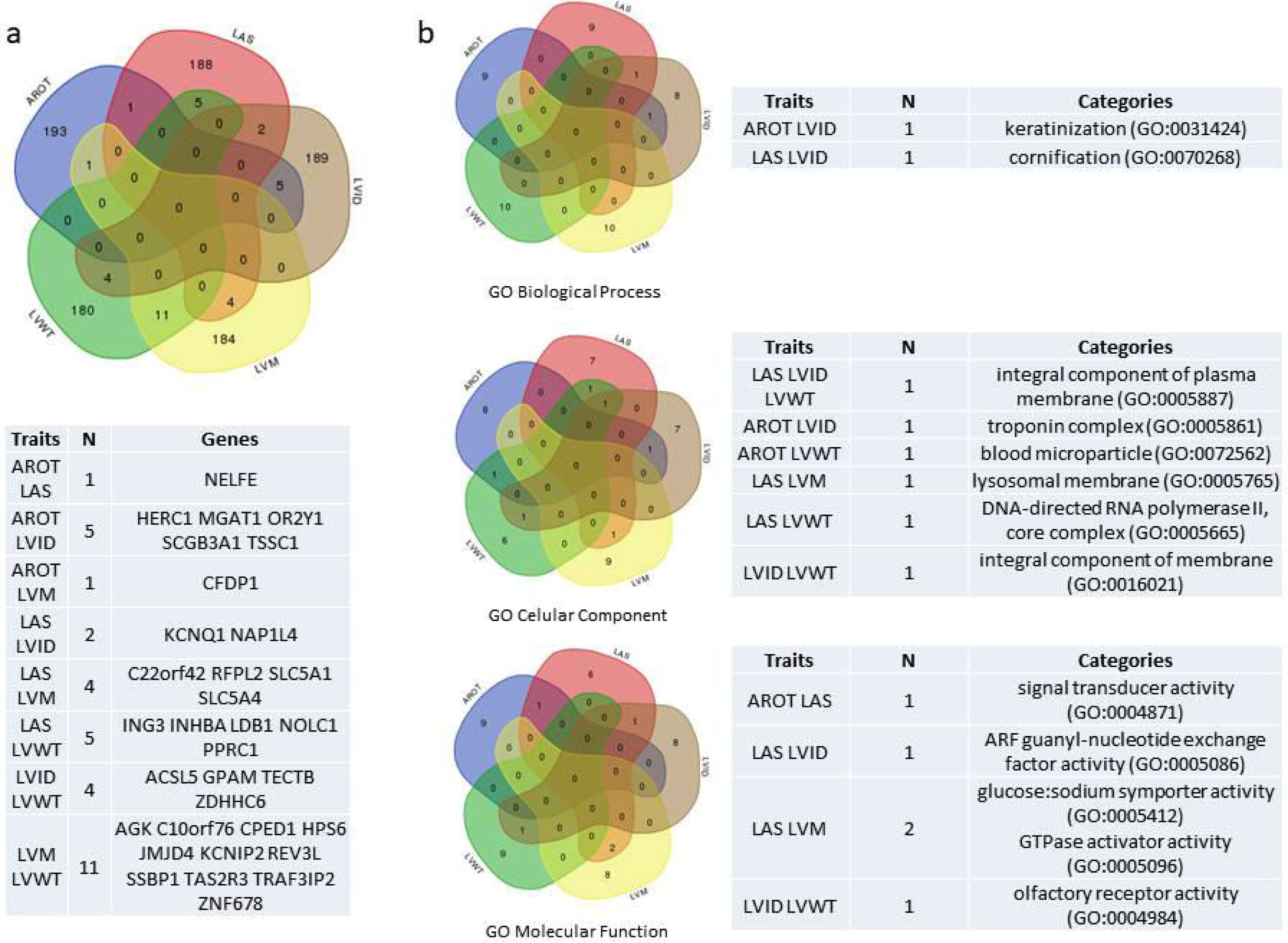
Venn diagrams showing the overlap of top genes (1a) and GO categories (1b) between the different traits analysed.

### Genetic correlation analysis between echocardiographic traits and AD

The association of gene variants related to echocardiographic measures with mental illnesses prompted us to explore in more depth the relationship between the echocardiographic phenotypes and AD. We performed a genetic correlation analysis using the study dataset (comprised by 11,559 individuals with echocardiographic phenotypes) along with 12,730 AD cases and controls both from internal and publicly available databases. First, we estimated the proportion of variance, as a proxy of trait heritability, explained by all SNPs in each one of these traits,^25^ which was higher for Alzheimer disease (0.38) than for the echocardiographic phenotypes (range: 0.17-0.36). Then, we looked for shared genetic loci between echocardiographic traits and AD using GREML analyses confirming the involvement of AD genes in heart performance detected during enrichment analyses. Specifically, we observed a positive correlation between AD and LAS (rG=0.167, p=0.0334), and negative correlations between AD and LVID (rG=-0.196, p=0.0056), and AD and LVM (rG=-0.198, p=0.0165); of note, LVM and LVID are the most correlated echocardiographic traits (rG=0.988, p<0.00001). The sign of the rG estimates determines whether a direct or inverse relationship between the two phenotype traits is observed. Therefore, our results suggest that SNPs that increase the risk of AD tend to be associated with increasing LAS values. On the contrary, we found that SNPs associated with increased risk of AD tend to be associated with decreasing ventricular measures (or *vice versa*), in particular LVID and LVM.

Based on these findings we performed a SNP-wise meta-analysis by pooling Fisher association p-values of two or more phenotypes. Thus, we combined in these meta-analyses p-values for LAS&AD, LVID&AD, LVM&AD, LAS&LVID&LVM&AD and LVID&LVM&AD and calculated gene-wise statistics (Supplementary tables 17-21). We performed a new enrichment analysis for identifying relevant functions and pathways determined by the top genes shared by AD and cardiac measures (Supplementary tables 22-25). The comparison of the enrichment results using Venn diagrams (Figure 3) showed that apoptosis related pathways driven by the *CASP8*, *CASP10* and *CFLAR* locus, and phospholipid scramblase genes are common elements for LAS, LVID, LVM and AD. We also observed an enrichment on genes present at the neuronal synapse such as glutamate and GABA receptors (*GRIN2C*, *GABRR1*, *GABRR2* and *GABBR2*), teneurin (*TENM2*), calsyntenin (*CLSTN3*), adenylate cyclase (*ADCY4*), *SLC5A7*, *LIN7A* or *LRFN2*. The haemoglobin complex, previously found among shared genes by LVID and LVM also seems to be relevant for AD.

**Figure 3.**
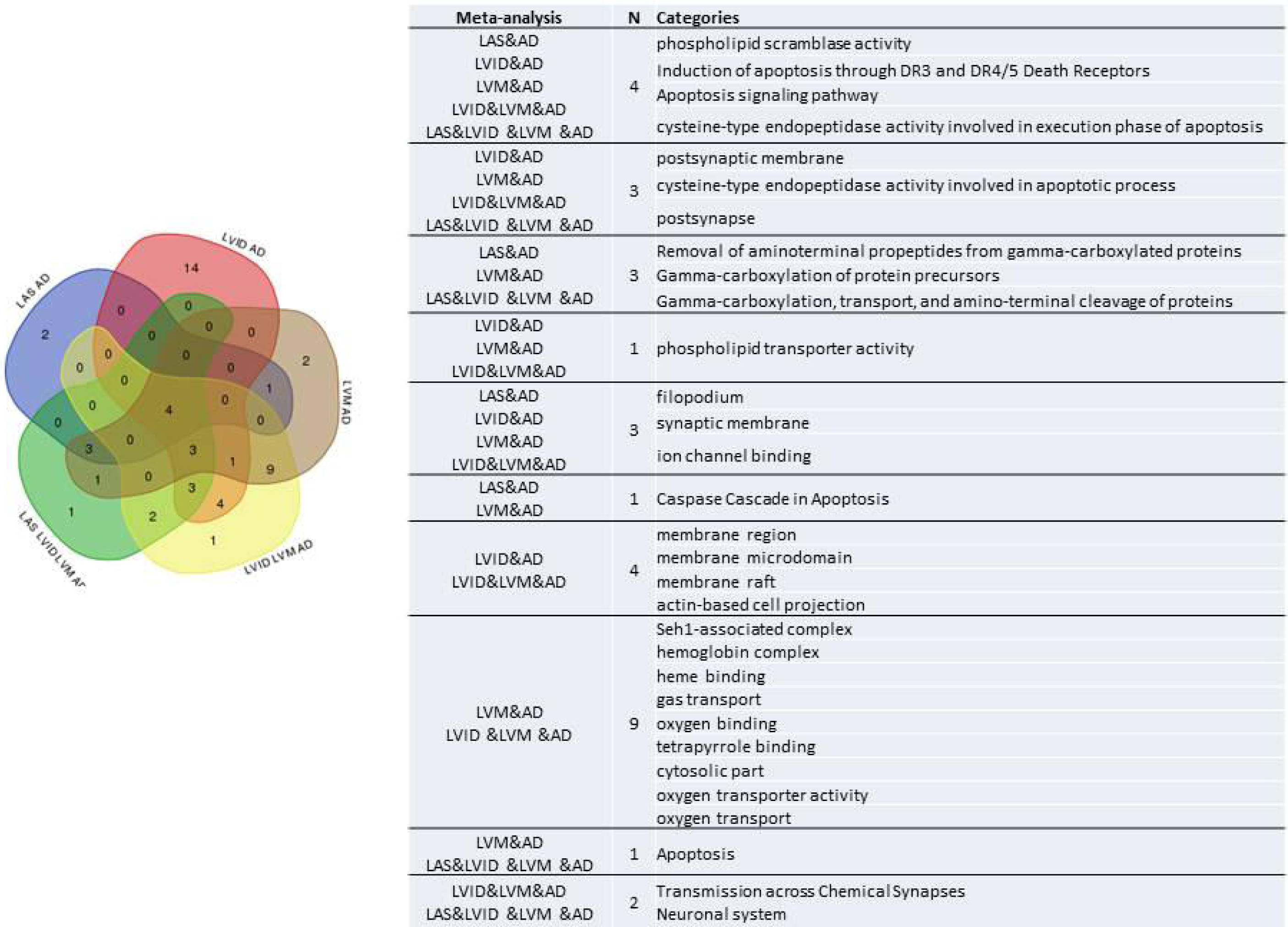
Enrichment analysis of top genes from meta-analysis on echocardiographic traits and Alzheimer’s disease.

## DISCUSSION

Our study, based on data from 11,559 individuals free of cardiovascular disease, shows that variants affecting diverse genes involved in apoptosis regulation associate with echocardiographic phenotypes in humans. We have obtained these results by both hypothesis-driven and agnostic approaches. In addition, novel findings from the analysis of our GWAS also include previously unnoticed associations of variants in genes involved in cell proliferation, DNA replication and mRNA splicing with left ventricular morphology. Our data also suggest the existence of a set of genes, mainly related to apoptosis/inflammation signalling, whose variants are associated with both cardiac phenotype and Alzheimer’s disease.

Our hypothesis-driven analysis showed that caspases 8 and 10 and the regulatory CFLAR (*cFLIP*) gene are strong predictors of LVM. Furthermore, our hypothesis-free approach found that caspase dependent pathways were overrepresented among the top 200 genes involved in the LVM phenotype. These results support our *a priori* hypothesis that the apoptotic signalling influences heart morphology with potential impact on heart performance. Our hypothesis was based in previous experimental work, including our own, showing that deficiency^8,10^ or overexpression^26^ of key apoptotic genes altered normal cardiomyocyte differentiation and heart development independently of cell death. Indeed, although these genes are best known for their role in regulating apoptotic cell death, experimental evidences show that the same genes also regulate myocyte proliferation, inflammation and hypertrophy in the heart.^8,10,27–29^ Because cell death is not a major event during heart development, and based on the above experimental work, we suggest that the relationship between the apoptotic genes and the cardiac phenotype might involve non-apoptotic functions.

In order to estimate the robustness of our GWAS analysis, we asked whether we had reproduced some signals already observed in previous genetic studies. Indeed, we found the already published link between *SMG6*, *TSR1* and *SRR* genes and AROT,^15,16^ but failed to detect statistically significant associations of other gene variants, possibly due to the limited power of this study. However, our analysis demonstrated genetic association between variants of genes previously shown to influence heart function and cardiac hypertrophy in experimental models, such as *MLF1,*^30^ which associates with LVM phenotype in our study, and *KCNIP2* (KChIP2)^31–33^ and *TRAF3IP2,*^34,35^ which have also been associated with LVM and LVWT phenotypes in humans in this study as well. Also, from the top list of genes whose variants are associated to LIVD, *ANKS6* has been associated with heart malformations.^36^

The GWAS analysis also showed strong association of variants of a group of 4 genes located in the 11p15.4 chromosomic region coding for odorant receptors with LVWT and LVID. Expression of these genes in non-neural tissues is related to the control of different processes, including glucose and oxygen homeostasis or cell cycle control,^37^ and has been shown to be involved in the regulation of cardiac function in rodent experimental models through interaction with fatty acids.^38^ Therefore, our genetic results open the possibility that odorant receptors’ activity can influence cardiac function in humans. Interestingly, low OR expression has been found in the cortex of neuro-psychiatric patients^39^ and a genetic microduplication in the 11p15.4 region has been associated with familial intellectual disability and autism.^40^

For the LVM phenotype, 8 of the 10 top ranked genes are involved in mRNA splicing, DNA replication and cell proliferation, and two are related to protein elongation and folding. Interestingly, 6 of the top genes associated with ventricular phenotypes have been also associated with mental illnesses including AD, Schizophrenia and bipolar disorder^41,42^ and the 11p15.4 region containing 4 of the 10 top genes for LVWT has been previously associated with intellectual disability.^40^

Unexpectedly, we found enrichment on AD related pathways for LAS, LVID and LVM that led us to explore comprehensively a possible link between AD and all these echocardiographic traits. Intriguingly, our results revealed for the first time a potential genetic link between AD and LAS, paired with a negative correlation between AD and either LVM or LVID. Interestingly, in line with our observation, a recent report found that LAS was independently associated to cognitive function in older adults.^43^ The opposite direction of the correlation coefficients for atrial and ventricular measures could be related to the different development patterns of these chambers during embryogenesis^44^ and with the described strong association of left atrial size with long-term exposure to vascular risk factors, particularly high blood pressure and obesity. In fact, the CARDIA Brain MRI Substudy found association of higher left atrial volume in early adulthood with impairment of white matter integrity in midlife, but not for ventricular measures.^45^

The link between AD and cardiac conditions is not well understood. The old concept of cardiogenic dementia was based on the high incidence of cardiac dysrhythmias observed in patients with dementia due to vascular causes.^46^ However, the relation between coronary heart disease (CHD) or HF and AD in epidemiological studies remains controversial, with some studies showing an association with cognitive impairment and dementia^47–49^ whereas others found no association.^50,51^ The fact that both conditions are competing risks complicates the study of their relationship. Lower cardiac index levels are related to lower cerebral blood flow in older adults free of CVD,^52^ but individuals with cardiac conditions that did not result in premature death might include many individuals chronically exposed to brain hypoperfusion due to reduced cardiac output that adaptively decreased cerebrovascular resistance through arteriolar dilatation. This kind of antagonistic pleiotropy between these phenotypes has been previously suggested by Beeri et al. after observing that better cognitive performance was associated with worse cardiac functioning in very elderly subjects.^53^

Moreover, whereas enlarged ventricular volume (LV hypertrophy) is a marker of diastolic dysfunction, LVM is also a marker of cardiovascular health, positively correlated with physical activity and cardiorespiratory fitness.^54,55^ Population based studies have shown an inverted U-shaped association for LVM values and age, since they rise in adolescence and decline with increased age.^56,57^ Furthermore, a U-shaped association between left ventricular ejection fraction (LVEF), a marker of systolic dysfunction, and abnormal cognitive decline has been reported, with increased dementia risk at the lowest and highest LVEF quintiles.^58^

To our knowledge, this is the first report analysing shared genetic factors between echocardiographic measures and AD. This method for estimating genome-wide pleiotropy has the advantage of being free of potential confounders determined by shared epidemiological risk factors such as high blood pressure or atherosclerosis. Our results show a negative genetic correlation for the ventricular measures LVID and LVM and AD, pointing to antagonist pleiotropic effects of shared genes by AD and LV cardiac measures, the main known functions of which are related to apoptosis, oxygen and phospholipid transport and neurotransmission. While low haemoglobin concentrations are associated with adverse cardiovascular outcomes and poor exercise capacity, increased concentration of the heme group have been found in the pathological lesions of AD patients.^59,60^ Hypoxia is a well known trigger factor for caspase related apoptosis, but caspases have been shown to play a role in cardiomyocyte morphogenesis rather than in cardiomyocyte death, whereas neuronal death is a hallmark of AD.^61^ Neurotransmission affects both cardiac and neuronal performance, and a few studies have examined synapse and neuron loss in AD brains and suggested that synaptic changes precede neuron loss.^62,63^ Finally, scramblase proteins, associated with LAS, LVID, LVM and AD in this study, have been involved not only in lipid transport, but also in mitochondrial membrane maintenance promoting cardiolipin synthesis, a key protein for myocardial energetics, and in caspase induced apoptosis.^64^ Anyway, lipid metabolism, especially cholesterol and fatty acid biogenesis have been associated with cardiovascular and cognitive phenotypes in adults being a potential common pathway explaining the connection between both groups of disorders and the decline of the incidence of dementia in younger cohorts with a better control of cardiovascular risk factors.^65^

In summary, our GWAS data suggest the influence of gene variants affecting the apoptotic/inflammation signalling pathway on left ventricular morphology and cardiac function, uncover novel candidate gene variants regulating echocardiographic phenotypes and establish a genetic link between cardiac morphology alterations, mental illness and Alzheimer’s disease involving key genes in the regulation of apoptotic signalling that deserve functional assessment due to their diagnostic and therapeutic potential.

## ACKNOWLEDGEMENTS

Data collection and sharing for this project was partially funded by the **Alzheimer’s Disease Neuroimaging Initiative (ADNI)** (National Institutes of Health Grant U01 AG024904) and DOD ADNI (Department of Defense award number W81XWH-12-2-0012). ADNI is funded by the National Institute on Aging, the National Institute of Biomedical Imaging and Bioengineering, and through generous contributions from the following: AbbVie, Alzheimer’s Association; Alzheimer’s Drug Discovery Foundation; Araclon Biotech; BioClinica, Inc.; Biogen; Bristol-Myers Squibb Company; CereSpir, Inc.; Cogstate; Eisai Inc.; Elan Pharmaceuticals, Inc.; Eli Lilly and Company; EuroImmun; F. Hoffmann-La Roche Ltd and its affiliated company Genentech, Inc.; Fujirebio; GE Healthcare; IXICO Ltd.; Janssen Alzheimer Immunotherapy Research & Development, LLC.; Johnson & Johnson Pharmaceutical Research & Development LLC.; Lumosity; Lundbeck; Merck & Co., Inc.; Meso Scale Diagnostics, LLC.; NeuroRx Research; Neurotrack Technologies; Novartis Pharmaceuticals Corporation; Pfizer Inc.; Piramal Imaging; Servier; Takeda Pharmaceutical Company; and Transition Therapeutics. The Canadian Institutes of Health Research is providing funds to support ADNI clinical sites in Canada. Private sector contributions are facilitated by the Foundation for the National Institutes of Health (www.fnih.org). The grantee organization is the Northern California Institute for Research and Education, and the study is coordinated by the Alzheimer’s Therapeutic Research Institute at the University of Southern California. ADNI data are disseminated by the Laboratory for Neuro Imaging at the University of Southern California.

The **AddNeuroMed** data are from a public-private partnership supported by EFPIA companies and SMEs as part of InnoMed (Innovative Medicines in Europe), an Integrated Project funded by the European Union of the Sixth Framework program priority FP6-2004-LIFESCIHEALTH-5. Clinical leads responsible for data collection are Iwona Kłoszewska (Lodz), Simon Lovestone (London), Patrizia Mecocci (Perugia), Hilkka Soininen (Kuopio), Magda Tsolaki (Thessaloniki), and Bruno Vellas (Toulouse), imaging leads are Andy Simmons (London), Lars-Olad Wahlund (Stockholm) and Christian Spenger (Zurich) and bioinformatics leads are Richard Dobson (London) and Stephen Newhouse (London). This dataset was downloaded from Synapse (doi:10.7303/syn2790911).

Funding support for the **Alzheimer’s Disease Genetics Consortium (ADGC)** was provided through the NIA Division of Neuroscience (U01-AG032984). This study was downloaded from NIH dbGaP repository (phs000372.v1).

The **Coronary Artery Risk Development in Young Adults Study (CARDIA)** is conducted and supported by the National Heart, Lung, and Blood Institute (NHLBI) in collaboration with the University of Alabama at Birmingham (N01-HC95095 & N01-HC48047), University of Minnesota (N01-HC48048), Northwestern University (N01-HC48049), and Kaiser Foundation Research Institute (N01-HC48050). This manuscript was not approved by CARDIA. The opinions and conclusions contained in this publication are solely those of the authors, and are not endorsed by CARDIA or the NHLBI and should not be assumed to reflect the opinions or conclusions of either. Genotyping for the CARDIA GENEVA cohort was supported by grant U01 HG004729 from the National Human Genome Research Institute. This study was downloaded from NIH dbGaP reporsitory ( phs000285.v3.p2).

The **Cardiovascular Heart Study (CHS)** was supported by contracts HHSN268201200036C, HHSN268200800007C, N01-HC85079, N01-HC-85080, N01-HC-85081, N01-HC-85082, N01-HC-85083, N01-HC-85084, N01-HC-85085, N01-HC-85086, N01-HC-35129, N01 HC-15103, N01 HC-55222, N01-HC-75150, N01-HC-45133, and N01-HC-85239; grant numbers U01 HL080295 and U01 HL130014 from the National Heart, Lung, and Blood Institute, and R01 AG-023629 from the National Institute on Aging, with additional contribution from the National Institute of Neurological Disorders and Stroke. A full list of principal CHS investigators and institutions can be found at https://chs-nhlbi.org/pi. This manuscript was not prepared in collaboration with CHS investigators and does not necessarily reflect the opinions or views of CHS or the NHLBI. Support for the genotyping through the CARe Study was provided by NHLBI Contract N01-HC-65226. This study was downloaded from NIH dbGaP reporsitory (phs000287.v5.p1).

The **Framingham Heart Study** is conducted and supported by the National Heart, Lung, and Blood Institute (NHLBI) in collaboration with Boston University (Contract No. N01-HC-25195 and HHSN268201500001I). This manuscript was not prepared in collaboration with investigators of the Framingham Heart Study and does not necessarily reflect the opinions or views of the Framingham Heart Study, Boston University, or NHLBI. “Funding for SHARe Affymetrix genotyping was provided by NHLBI Contract N02-HL64278. SHARe Illumina genotyping was provided under an agreement between Illumina and Boston University. Funding for Affymetrix genotyping of the FHS Omni cohorts was provided by Intramural NHLBI funds from Andrew D. Johnson and Christopher J. O’Donnell. This dataset was obtained from the NIH dbGaP repository (phs000007.v29.p10).

The genotypic and associated phenotypic data used in the study, “Multi-Site Collaborative Study for Genotype-Phenotype Associations in Alzheimer’s Disease (**GenADA**)” were provided by the GlaxoSmithKline, R&D Limited. The datasets used for analyses described in this manuscript were obtained from NIH dbGaP repository (phs000219.v1.p1).

The **Mayo** Clinic Alzheimer’s Disease Genetic Studies, led by Dr. Nilüfer Ertekin-Taner and Dr. Steven G. Younkin, Mayo Clinic, Jacksonville, FL using samples from the Mayo Clinic Study of Aging, the Mayo Clinic Alzheimer’s Disease Research Center, and the Mayo Clinic Brain Bank. Data collection was supported through funding by NIA grants P50 AG016574, R01 AG032990, U01 AG046139, R01 AG018023, U01 AG006576, U01 AG006786, R01 AG025711, R01 AG017216, R01 AG003949, NINDS grant R01 NS080820, CurePSP Foundation, and support from Mayo Foundation. This dataset was downloaded from Synapse (doi:10.7303/syn5550404).

The **MESA study** was supported by contracts HHSN268201500003I, N01-HC-95159, N01-HC-95160, N01-HC-95161, N01-HC-95162, N01-HC-95163, N01-HC-95164, N01-HC-95165, N01-HC-95166, N01-HC-95167, N01-HC-95168 and N01-HC-95169 from the National Heart, Lung, and Blood Institute, and by grants UL1-TR-000040, UL1-TR-001079, and UL1-TR-001420 from NCATS. The authors thank the other investigators, the staff, and the participants of the MESA study for their valuable contributions. A full list of participating MESA investigators and institutions can be found at http://www.mesa-nhlbi.org. This dataset was obtained from the NIH dbGaP repository (phs000209.v6.p2).

The **Neocodex-Murcia** study was funded by the Fundación Alzheimur (Murcia), the Ministerio de Educación y Ciencia (Gobierno de España), Corporación Tecnológica de Andalucía and Agencia IDEA (Consejería de Innovación, Junta de Andalucía). The Diabetes Research Laboratory, Biomedical Research Foundation. University Hospital Clínico San Carlos has been supported by CIBER de *Diabetes y Enfermedades Metabólicas Asociadas* (CIBERDEM); CIBERDEM is an ISCIII Project.

The **ROS/MAP** study data were provided by the Rush Alzheimer’s Disease Center, Rush University Medical Center, Chicago. Data collection was supported through funding by NIA grants P30AG10161, R01AG15819, R01AG17917, R01AG30146, R01AG36836, U01AG32984, U01AG46152, the Illinois Department of Public Health, and the Translational Genomics Research Institute. This dataset was downloaded from Synapse (doi:10.7303/syn3219045).

The **TGEN** study was supported by Kronos Life Sciences Laboratories, the National Institute on Aging (Arizona Alzheimer’s Disease Center P30 AG19610, RO1 AG023193, Mayo Clinic Alzheimer’s Disease Center P50 AG16574, and Intramural Research Program), the National Alzheimer’s Coordinating Center (U01 AG016976), and the state of Arizona. TGEN investigators provided free access to genotype data to other researchers via Coriell Biorepositories (http://www.coriell.org).

The results published here are in part based on data obtained from the AMP-AD Knowledge Portal accessed at doi:10.7303/syn2580853.

## ETHICS STATEMENT

This study complies with the Declaration of Helsinki and a locally appointed ethics committee has approved the research protocol.

## APPENDIX: ABBREVIATIONS

LVM: Left Ventricular (LV) Mass (g)
LVID: End-Diastolic LV Internal Dimension (cm)
LVWT: LV Wall Thickness (cm) (TPW+TIS)
LAS: Left Atrial Size (cm)
AROT: End-Diastolic Diameter of the Aortic Root (cm)
LV: Left Ventricle
CVD: Cardiovascular Disease
CHD: Coronary Heart Disease
HF: Heart Failure
TPW: End-Diastolic Thicknesses of the Posterior Wall
TIS: End-Diastolic Thicknesses of the Interventricular Septum

## ADDITIONAL INFORMATION

AR and DS contributed to the design of the study, acquisition of data and interpretation of the results; MES contributed to the design, acquisition of data, statistical analysis, interpretation of the results and redaction of the manuscript; AG contributed to the design of the study and interpretation of the results and redaction of the manuscript; BH-O, SMG, IR and AB contributed to the statistical analysis and critical review of the manuscript; GM, AO and SV contributed to the acquisition of data and critical review of the manuscript; JXC critically reviewed the manuscript.

